# ICeD-T Provides Accurate Estimates of Immune Cell Abundance in Tumor Samples by Allowing for Aberrant Gene Expression Patterns

**DOI:** 10.1101/326421

**Authors:** Douglas R. Wilson, Joseph G. Ibrahim, Wei Sun

**Affiliations:** University of North Carolina at Chapel Hill; Fred Hutchinson Cancer Research Center

**Keywords:** Immunotherapy, Immuno-Oncology, Bulk Expression, Deconvolution, RNA-seq, Microarray

## Abstract

Immunotherapies have achieved phenomenal success in the treatment of cancer and promise even more breakthroughs in the near future. The need to understand the underlying mechanisms of immunotherapies and to develop precision immunotherapy regimens has spurred great interest in characterizing immune cell composition within the tumor microenvironment. Several methods have been developed to estimate immune cell composition using gene expression data from bulk tumor samples. However, these methods are not flexible enough to handle aberrant patterns of gene expression data, e.g., inconsistent cell type-specific gene expression between purified reference samples and this cell type in tumor samples. In this paper, we present a novel statistical model for expression deconvolution called ICeD-T (Immune Cell Deconvolution in Tumor tissues), which models gene expression by a log-normal distribution that is appropriate for both microarray and RNA-seq data. ICeD-T automatically identifies aberrant genes whose expressions are inconsistent with the deconvolution model and down-weights their contributions to cell type abundance estimates. We evaluated the performance of ICeD-T versus existing methods in simulation studies and several real data analyses. ICeD-T displayed comparable or superior performance to these competing methods. Applying these methods to assess the relationship between immunotherapy response and immune cell composition, ICeD-T is able to identify significant associations that are missed by its competitors.

## 1 Introduction

The evolving relationship between a cancer and its host’s immune system is well summarized by a hypothesis known as immunoediting. Immunoediting stresses that the immune system not only suppresses tumor cells, but also shapes tumor immunogenicity in ways that may promote tumor growth [1, 2]. For example, consider the relationship between tumors and tumor-infiltrating T cells. Infiltrating T cells can be cytotoxic, contributing to the reduction of cancer cell populations. However, these T cells also express immune checkpoints that inhibit their function, and such checkpoints can prevent the immune system from indiscriminately attacking healthy host cells. Under selective pressure from the immune system, cancers cells may exploit the immune checkpoints to escape the attack by infiltrating T cells.

Early strategies in immunotherapy were developed based on the insights of immunoediting [3]. Among the best known immunotherapy strategies, immune checkpoint inhibitors block immune inhibition pathways that restrict effective anti-tumor T cell responses [4]. Checkpoint inhibitors have achieved phenomenal success in a fraction of cancer patients, exhibiting response rates around 40% and 20% for melanoma and lung cancer, respectively
[5]. It is of great clinical interest to identify the subset of cancer patients who may respond to checkpoint inhibitors. Use of tumor-infiltrating immune cells to predict clinical response to therapy has shown promising results. Previous studies have shown that the patients with CD8+ T cells around tumor cells have higher response rate to checkpoint inhibitors
[6]. In addition to benefiting the development of precision immunotherapies, immune cell composition in tumor samples have also demonstrated prognostic value [7, 8]. Therefore, studying immune cell composition in tumor samples is timely and potentially has high impact on cancer research.

Several groups have studied immune cell composition using gene expression data from bulk tumor samples [9–14]. These pioneering works have demonstrated promising results, but also bear some limitations. For example, a subset of these works estimate immune cell presence using the expression of few genes [9, 10], or calculate average expression of the genes with cell type-specific expression [15] instead of estimating immune cell composition. As an alternative, several methods have been proposed to estimate immune cell composition using a regression-based approach, with gene expression from bulk tumor samples as the response variable and reference gene expression from purified cell types as covariates. For example, CIBERSORT [12] employs support-vector regression. TIMER [13] uses a linear regression and removes the genes with very high expression due to their strong influence on model fitting. EPIC [14] is the most recent work. It uses weighted linear regression to give the genes with lower expression variation higher weights. These regression-based methods, when applied to tumor expression data, explicitly or implicitly assume that they start with a set of genes that have negligible expression in tumor cells, and that the expression of immune cells are conserved between purified reference samples and tumor samples. These assumptions are questionable as many environmental factors that affect gene expression may differ between tumor and reference samples.

In this paper, we propose a new statistical method for cell type deconvolution entitled ICeD-T, which stands for Immune Cell Deconvolution in Tumor tissues. ICeD-T is an extension of existing regression based methods [12–14] with two major novel features designed to overcome the limitations of these methods.

First, ICeD-T employs a likelihood based framework, which assumes that gene expression follows a log-normal distribution. Previous work has shown that deconvolution should be performed on linear-scale instead of log-scale of gene expression data since linear-scale mixing of gene expression better captures the biological reality [16]. However, since gene expression variation increases with expression level, genes with higher expression may become outliers with great influence on linear scale deconvolution. Therefore one may need to remove genes with high expression for robust deconvolution analysis [13]. The log transformation, often used in expression studies, enjoys variance-stabilizing and skew-mitigation properties that limit the impact of genes with higher expression [17, 18]. ICeD-T is able to perform gene expression deconvolution on the linear-scale while simultaneously incorporating the beneficial properties of the log-transformation through our method design and the use of log-normal distribution.

Second, ICeD-T automatically identifies the genes whose expression in tumor samples are inconsistent with reference profiles due to altered expression in tumor infiltrating immune cells or unexpected tumor cell expression. Within its estimation algorithm, ICeD-T down-weights the contribution of such genes in cell type abundance estimation using a mixture model that separates all the genes into two groups: an “aberrant” group and a “consistent” group.

## 2 Statistical Methods

### 2.1 The Input Data

While ICeD-T can be applied on microarray data, we focus mainly on RNA-seq data as it is more popular now and in the foreseeable future. We assume that RNA-seq data from bulk tumor samples are available for *n* independent subjects. Gene expression from purified samples may be pre-computed or processed from raw RNA-seq data of multiple replicates for each cell type. Across reference expression profiles and bulk samples, the RNA-seq measurements of gene expression are appropriately normalized in a consistent manner using FPKM, FPKM-UQ, or TPM. More specifically, to calculate FPKM, we divide gene expression (# of RNA-seq fragments) by total number of mapped fragments per sample (in millions) and the gene length (in kilo bases). FPKM-UQ is a variant of FPKM where sample-specific read-depth is measured by 75 percentile of gene level fragment counts across all genes, instead of the total number of mapped fragments. TPM reverses the order of the two normalization steps. It first divides the gene-level fragment counts by gene length, and then divides it by the summation of gene-length corrected fragment counts across all genes.

Additional information utilized by ICeD-T’s deconvolution model includes a pre-selected gene set (ideally, genes with immune-specific expression) and tumor purity, if available. Several such gene sets have been prepared by previous work, such as the gene sets used by CIBERSORT or EPIC [12, 14]. Provision of tumor purity is optional, and it can be computed, for example, using somatic copy number alteration data [19].

### 2.2 Statistical Model

Specification of the ICeD-T model begins with a consideration of expression behavior in purified references samples of constituent cell types. Denote by *Z*_*jkh*_ the expression of gene *j* in the *h*-th purified sample of cell type *k*. ICeD-T assumes that the *Z*_*jkh*_’s follow independent log-normal distributions, given by:

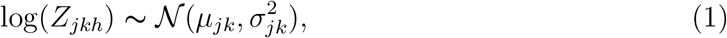

where

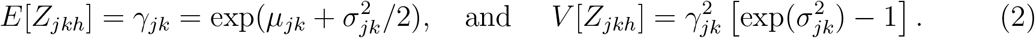

Therefore, the distribution parameters for each cell type’s gene expression (e.g.,*μ*_*jk*_ and 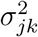) may be estimated by the mean and variance of the log-transformed *Z*_*jkh*_ values. Once estimated, these parameters represent expression profiles for each cell type in our deconvolution model. Optionally, ICeD-T accepts previously computed profiles which would replace the *γ*_*jk*_ above.

Shift focus to the *n* bulk tumor samples. Assuming that each sample is composed of *K* immune cell types and other extraneous cell types, the expression of gene *j* in bulk tumor sample *i* - denoted by *Y*_*ij*_ - is modeled by

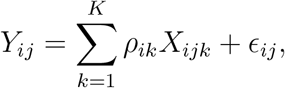

where *X*_*ijk*_ represents the expression of gene *j* for cells of type *k* in the *i*-th sample, and *ρ*_*ik*_ is the proportion of expression attributable to cell type *k*. The residual error *ϵ*_*ij*_ represents signals from other cell types (e.g., tumor cells) or random noise. If tumor purity information is provided, 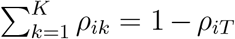, where *ρ*_*iT*_ is tumor purity. If tumor purity is not provided,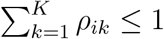.

We begin to develop the probabilistic framework utilized by ICeD-T to model the relationship posited above by first assuming that there are no aberrant genes (i.e., gene expression of each cell type in reference samples is consistent with gene expression in tumor microenvironment). Under such an assumption, *X*_*ijk*_ has the same distribution as the *Z*_*jkh*_ for any *i*, *h*, and *j* (i.e., *X*_*ijk*_ ~ *Z*_*jkh*_). The summation of independent log-normal random variables does not have a closed form distribution function. To address this issue, ICeD-T approximates the distribution of *Y*_*ij*_ using another log-normal:

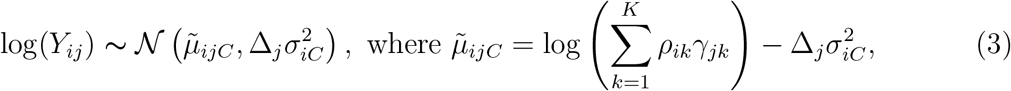
and Δ_*j*_ is the weight for the *j*-th gene.

The approximation used above is based upon the Fenton-Wilkinson approach which states that the summation of log-normals can be approximated by another log-normal whose parameters are obtained via moment-matching [20]. Under a strict Fenton-Wilkinson approach, the distribution of *Y*_*ij*_ would be given by:

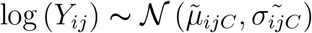

where

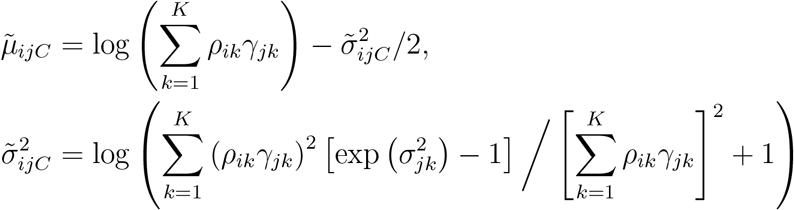

We replace the variance structure posited by Fenton-Wilkinson with the weighted variance model of equation (3) as the weighted model demonstrated improved fit and stability in simulated data.

Regarding the variance weights used by ICeD-T, we implement two different options. One assumes a homogeneous weight for all genes, i.e., Δ_*j*_ = 1 for all *j*. Later we refer to this option as “No Weights”. The other weight option is termed maximal variance weights or “Max Var Weights”. To define maximal variance weights, let 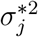 be the maximum expression variance across all cell types *k* at gene *j*:

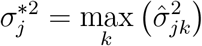

The weight of a gene *j* is then specified as follows:

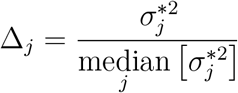

Thus, a gene’s weight compares its maximal expression variance to the median of all such maxima across genes. Under this construction, genes with larger variances will have larger variance weights. Larger variance weights ensure that residuals from such genes will have smaller impact on estimation of cell type composition.

The Δ*j* specified above requires slight modification to improve stability of the model fit. Unadjusted, this procedure can provide some genes with excessively small variance weights and some genes with excessively high variance weights. To control this extreme behavior, the bottom 15% of variance weights are replaced with the 15th percentile variance weight across all genes. Similarly, the top 15% of all variance weights are replaced by the 85th percentile variance weight. In this way, no genes are allowed to become too minimally or maximally important to model fit.

Return to the specification of *Y*_*ij*_ in equation (3). Now assume that some genes in the dataset are aberrant. For aberrant genes, ICeD-T borrows the expression structure proposed for consistent genes but inflates the variance. Thus, if gene *j* is aberrant, the expression of *Y*_*ij*_ is given by:

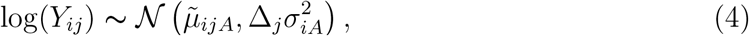

where

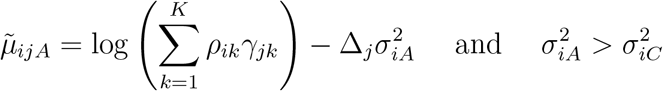

By allowing aberrant genes to have larger variance, the ICeD-T model flattens the likelihood for such genes, and thus down-weights their contributions to cell type proportion estimates.

Direct use of the likelihoods provided by equations (3) and (4) within bulk data is impossible since it is unknown whether a gene is consistent or aberrant *a priori.* Thus, ICeD-T must model expression at any gene as a mixture of the log-normal distributions pertaining to consistent and aberrant genes. The mixture likelihood utilized by ICeD-T is found below:

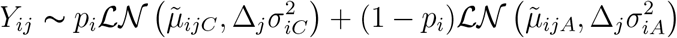

where 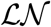 denotes the density function of a log-normal distribution, and *p*_*i*_ and 1 – *p*_*i*_ denotes the proportion of genes being consistent and inconsistent, respectively. This likelihood function can be maximized using an EM algorithm. Missing data necessary for the EM algorithm is introduced in the form of class membership indicators *H*_*ij*_, where *H*_*ij*_ = 0 or 1 denotes whether the *j*-gene is aberrant or consistent in the *i*-th bulk tumor sample, respectively. Thus, the complete data log-likelihood for the *i*-th bulk tumor sample is given by:

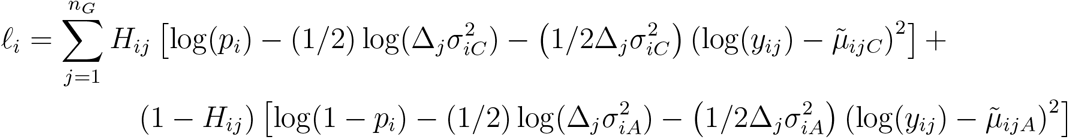

where *n*_*G*_ is the number of genes used in our model.

Within each EM step, maximization of *Q* function with respect to 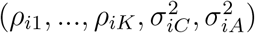 and *p*_*i*_ are separable. Given the other parameters, the estimate of *p*_*i*_ has a closed form. Given *p*_*i*_, the remaining parameters are grouped into two blocks: the mixture proportions *ρ*_*ik*_’s (block 1) and the two variance parameters 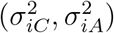 (block 2). The parameters of these two blocks are iteratively updated. Given the estimates of 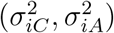, the mixture proportions *ρ*_*ik*_ are estimated using numerical optimization (the BFGS algorithm) while the constraints are incorporated using the Augmented Lagrangian method (R function **`auglag`**). Given the estimates of the mixture proportions *ρ*_*ik*_’s, the two variance terms 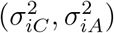 are involved in separate pieces of the complete data log-likelihood, and thus can be estimated separately. Given variance weights, each of 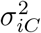 and 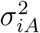 is estimated by numerical optimization (R function **`optimize`**). Without variance weights, they can be estimated by closed form. See Supplementary Materials Section A for details of the parameter estimation steps.

The *ρ*_*ik*_’s estimated by any regression based deconvolution approach should be interpreted as the proportion of gene expression contributed by certain cell types. If one seeks to estimate the proportion of cells, these *ρ*_*ik*_’s should be adjusted by cell size factors. We borrow the cell size factors, denoted by *s*_*k*_, from Racle et al. [14] and construct revised relative abundance of immune cell types by 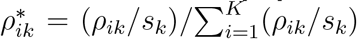 Further details are provided in the Supplementary Materials (Section C.2).

### 3 Results

#### 3.1 Simulation Study

We conducted a simulation study to evaluate the performance of ICeD-T, CIBERSORT, and EPIC. For each method, we seek to assess the estimation accuracy and the robustness of estimation in the presence of aberrant gene behavior. For ICeD-T only, we also assess its ability to identify aberrant genes.

We simulated reference expression of 250 genes for 5 cell types: one tumor cell type and four immune cell types. Our simulations assume that these 250 genes were selected to be expressed in immune cells but not tumor cells. When there are no aberrant genes, the expression of these 250 genes in a bulk tumor sample was simulated by mixing the 4 immune cell types with known proportions. For each gene, we assume it is expressed in one of the four immune cell types and has low/background expression in the other three immune cell types. To better mimic the complexity of real data, we do not assume one homogeneous background expression. Instead, we assume the background expression has a three-tiered scale to reflect lowly, moderately or highly expressed genes (range in log scale: 2.0-8.0). Average log-transformed expression for the expressed cell type is simulated from by an up-shift of background expression level (range: 3.5-9.0). See Supplementary Materials Section B.1 for more details. Using RNA-seq expression data from immune cells taken from Linsley et al. [21], a mean-variance relationship was computed from FPKM-UQ normalized data across immune specific genes. Then we used this mean-variance relationship and the simulated average expression profiles to decide corresponding variance with allowance for random error. Fifteen reference samples were simulated for each cell type from its unique expression profile using a log-normal distribution.

To generate the expression of a bulk tumor sample, a tumor purity value was simulated from a normal distribution (mean=0.60, sd=0.15) and truncated at endpoints of 0.17 and 0.95. The remaining immune cell proportions were then simulated from a Dirichlet distribution with average abundances ranging from 15% to 40%. For each gene in the bulk tumor sample, its expression in each immune cell type was simulated from a log-normal distribution and a weighted summation of these expression values was computed as the expression in the bulk tumor sample. These gene expression profiles are then perturbed to account for aberrant behavior. Zero or approximately twenty percent of genes were randomly selected as aberrant genes. Among them, 25% have down-regulated expression in the highly expressed cell type, 25% have up-regulated expression of the highly expressed cell type, and 50% have expression in tumor cells at a background level. See the Appendix B for further details regarding the construction of these simulations and additional simulation results.

The expression profile of each cell type was estimated from the 15 simulated samples of that cell type. This reference is used for deconvolution in each of the following models: ICeD-T without variance weights, ICeD-T with variance weights, LNorm with variance weights, CIBERSORT (version Jar 1.06), and EPIC. LNorm stands for “log normal”, and it is a variant of the ICeD-T model which does not consider aberrant gene behavior.

When there is no aberrance in gene expression, all methods perform well, while ICeD-T provides the most accurate estimates of cell type proportions (Figure 1). When 20% of the 250 genes are aberrant, the performance of LNorm, EPIC, and CIBERSORT all become worse, while the performance of ICeD-T method remain similar (Figure 2). Both EPIC and LNorm’s cell type proportion estimates suffer from bias and larger variance in the presence of aberrant genes. CIBERSORT still performs relatively well, but has an apparent inflation of the estimation variance. While the weighted version of ICeD-T provides the best results, both weighted and unweighted ICeD-T are able to maintain high accuracy with minimal estimation variance (Figure 2(a)-(b)).

**Figure. 1.**
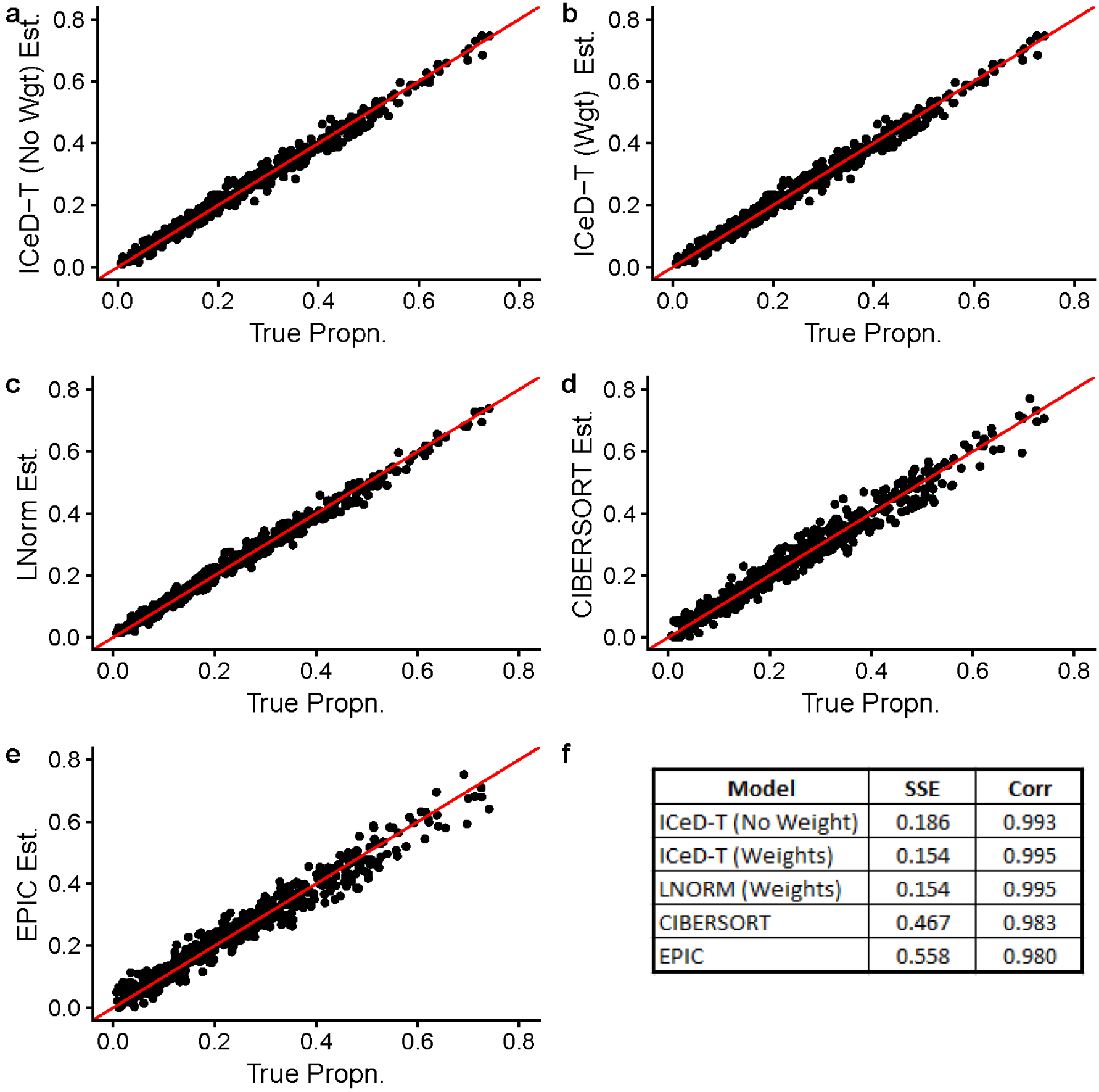
Visualized results of model fits on simulated data without aberrance. Figure (f) summarizes the accuracy across all 135 subjects for each model.

**Figure. 2.**
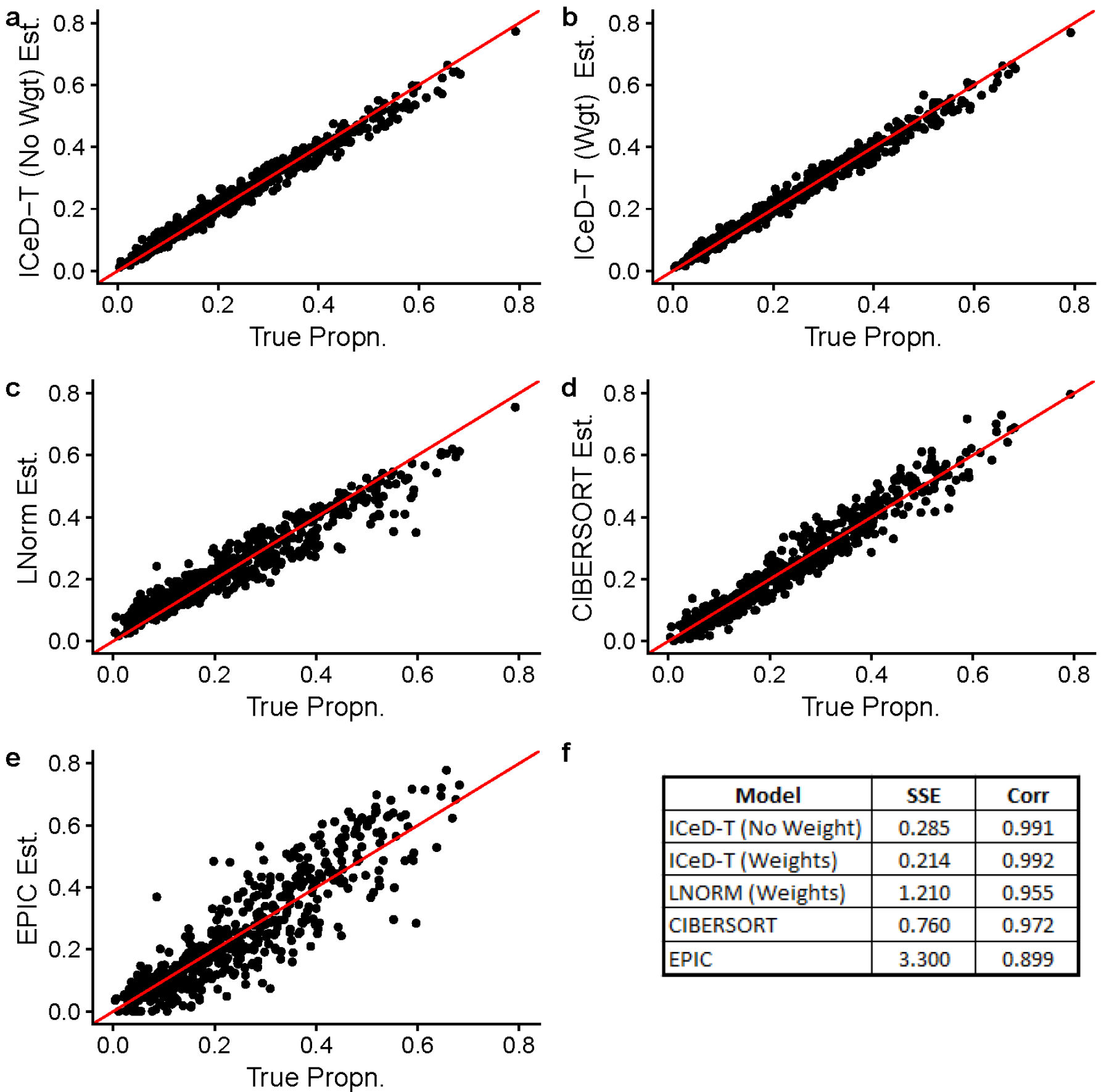
Visualized results of model fits on simulated data when ~ 20% of the genes are abberant. Figure (f) summarizes the accuracy across all 135 subjects for each model.

**Figure. 3.**
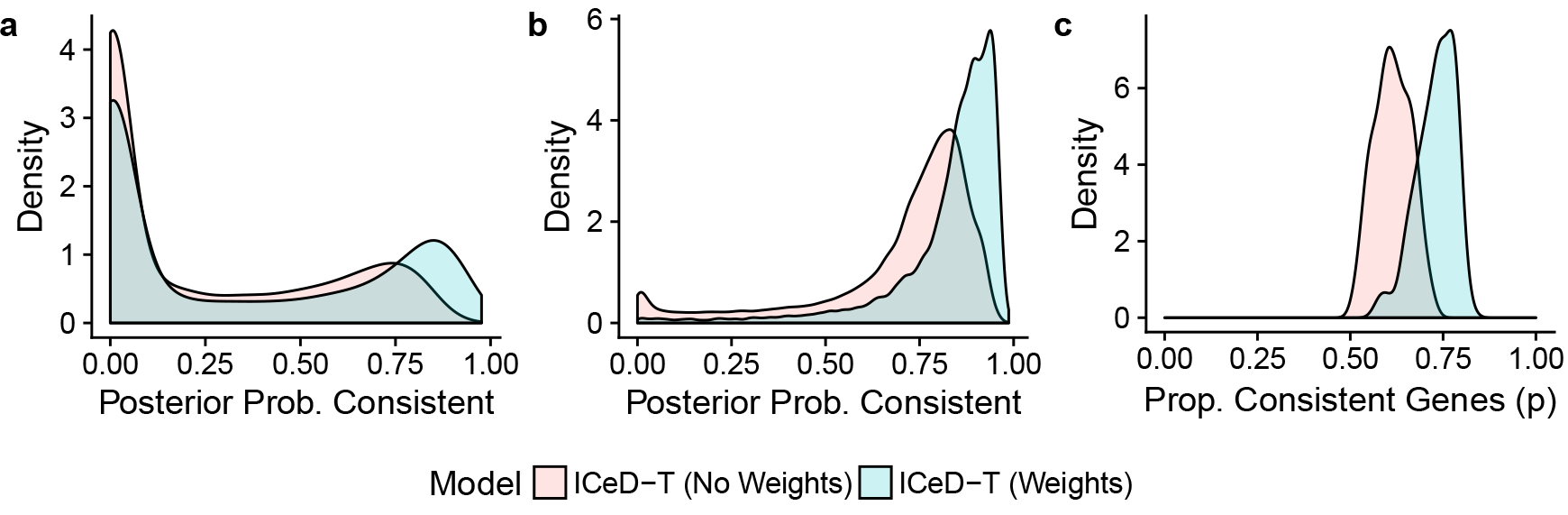
(a) The posterior probabilities of being consistent for those aberrant genes. (b) The posterior probabilities of being consistent for those consistent genes. (c) Estimates of the proportion of consistent genes.

To identify aberrant genes, ICeD-T computes the posterior probability of a gene being consistent. Examining the distribution of this quantity across consistent and aberrant genes, we see that both the weighted and unweighted versions of ICeD-T separate consistent and aberrant genes reasonably well (Figure 3). The weighted variant of ICeD-T provides more accurate estimate of the proportion of aberrant genes, and identify consistent genes with higher confidence. For aberrant genes, the posterior probability of being consistent show a bi-modal distribution, implying that a small proportion of aberrant genes are missed. This is partly due to our very challenging simulation setting, with three types of aberrant patterns and three tiers of expression levels for background genes. Such three tiers of background diminishes the difference between background cell types and expressed cell types, and further complicates the identification of aberrant genes.

#### 3.2 Validation in Microarray Expression of PBMCs

In the CIBERSORT paper, Newman et al. [12] described the collection of peripheral blood mononuclear cell (PBMC) gene expression data from 20 healthy adults. After extraction of PBMC samples from each subject, these samples were subjected to microarray expression analysis and flow cytometric measurement to establish ground-truth of cell type proportions. We use this dataset to evaluate our method and compare its performance with CIBERSORT and EPIC.

To be consistent with the approach used by Newman et al. [12], we use the their LM22 reference of cell type-specific gene expression for all methods. The LM22 reference matrix is derived from microarray gene expression data, and thus is consistent with the gene expression platform of the bulk tissue samples. EPIC had developed its own reference matrices from RNA-seq data (TRef for bulk tumor samples and BRef for bulk normal samples), but they are inappropriate in microarray settings. Because EPIC and ICeD-T both require that the gene expression from bulk samples and reference samples are measured on the same scale, gene expression data from bulk samples were quantile normalized to a target distribution established by the reference samples used to derive the LM22 matrix. The results of each method are then restricted to the nine cell-types examined in Newman et al. [12]: naive B-cells, memory B-cells, CD8+ T-cells, naive/memory resting/memory activated CD4+ T-cells, *γδ* T-cells, Natural killer cells, and monocytes. Estimates for each mixture sample are renormalized so that their summation equals 100 after correction for cell size of different cell types. The accuracy of each method is assessed by comparing sums of squared errors and correlations between the expression-based cell type proportion estimates and flow-cytometry estimates. Correlations are computed by pooling cell type proportions for all subjects and all cell types.

**Table 1:** Validation of immune cell proportion estimates by flow cytometry for 9 cell types [left] and 6 cell types after grouping naive B-cells and memory B-cells as B cells, and naive/memory resting/memory activated CD4+ T-cells as CD4+ T cells [right].

| Model | SSE | Cor | Model | SSE | Cor |
| --- | --- | --- | --- | --- | --- |
| ICeD-T (no weight) | 13.10 | 0.53 | ICeD-T (no weight) | 10.48 | 0.75 |
| ICeD-T (w/ weight) | 12.05 | 0.59 | ICeD-T (w/ weight) | 9.44 | 0.78 |
| CIBERSORT | 14.15 | 0.65 | CIBERSORT | 11.02 | 0.77 |
| EPIC | 29.43 | 0.31 | EPIC | 32.01 | 0.18 |

**Figure 4:**
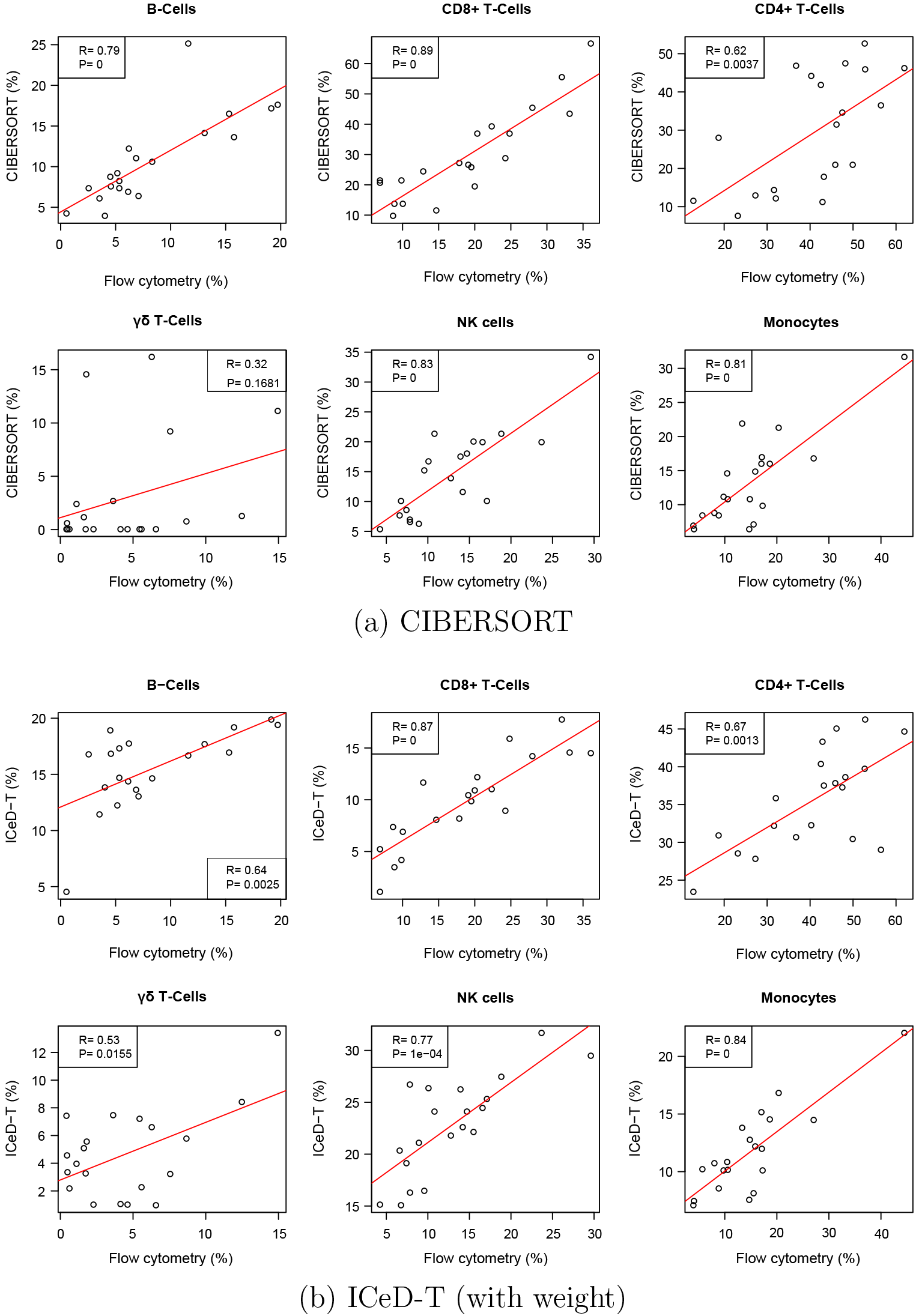
Comparison of cell type proportion estimates by CIBERSORT and ICeD-T versus the cell type proportions measured by flow cytometry. Red lines indicate the least squares model fit to the estimated immune proportions.

Examining the results of the 9 original cell types, ICeD-T provides the most accurate estimates of cell type proportions in terms of sum of squared errors. CIBERSORT, on the other hand, provides the most accurate estimates with respect to the correlations (Table 1, Figure 4). However, the superior correlation of CIBERSORT is due in part to several cell subsets with positive correlations but severe bias (e.g., memory activated CD4 T-cells, memory resting CD4 T-cells) (Supplementary Materials Section C.4). After grouping a few highly similar cell types (e.g., grouping naive B-cells and memory B-cells as B cells, and naive/memory resting/memory activated CD4+ T-cells as CD4+ T cells), ICeD-T achieves comparable or higher correlation between expression-based cell type proportion estimates and flow-cytometry estimates while maintaining the smallest sum of squared errors (Table 1, Figure 4). In this dataset, EPIC has very poor performance, which may be due to the fact that it is designed for RNA-seq data.

#### 3.3 Flow Cytometry Validation in Melanomas

In the EPIC paper, Racle et al. [14] obtained metastatic melanoma samples from the lymph nodes of four patients with stage III melanomas. A portion of each of these samples was used for a flow cytometric analysis while the remaining portion was used for bulk RNA-sequencing. Results from flow cytometry were used to establish a ground-truth cell type composition. TPM-normalized RNA-seq expressions and flow cytometry measured compositions were extracted directly from the EPIC R package.

We used EPIC’s TRef matrix as reference gene expression for both EPIC and ICeD-T. ICeD-T was run in four different modes, with or without variance weights (denoted by wY and wN, respectively) and with or without sample purity as part of the inputs (denoted by pY and pN, respectively). For this analysis, purity is defined as the proportion of non-immune content plus the proportions of cells not assessed via flow cytometry (e.g. macrophages, fibroblasts, and endothelials, and others). CIBERSORT was fit using both the LM22 and TRef matrices directly to the TPM data. All cell type proportion estimates were corrected by cell size factors reported by Racle et al. [14]. To allow comparison of ICeD-T and EPIC with CIBERSORT that only computes relative immune cell abundance estimates, we obtain relative proportions for all methods by normalizing cell type proportions so that they add up to 1.

Overall EPIC provides more accurate estimates of the total proportion of all immune cells, while ICeD-T provides more accurate estimation of the relative proportions of immune cells among the modeled immune cell types (Table 2, Figure 5, Supplementary Materials, Section D.3). Comparing non-relativized proportions of the remaining immune cells, ICeD-T (pY, wY) improves upon EPIC’s fit in terms of the overall sum of squared error (0.043 vs 0.11) while preserving strong correlation (0.924 vs 0.918) across all subjects.

**Figure 5:**
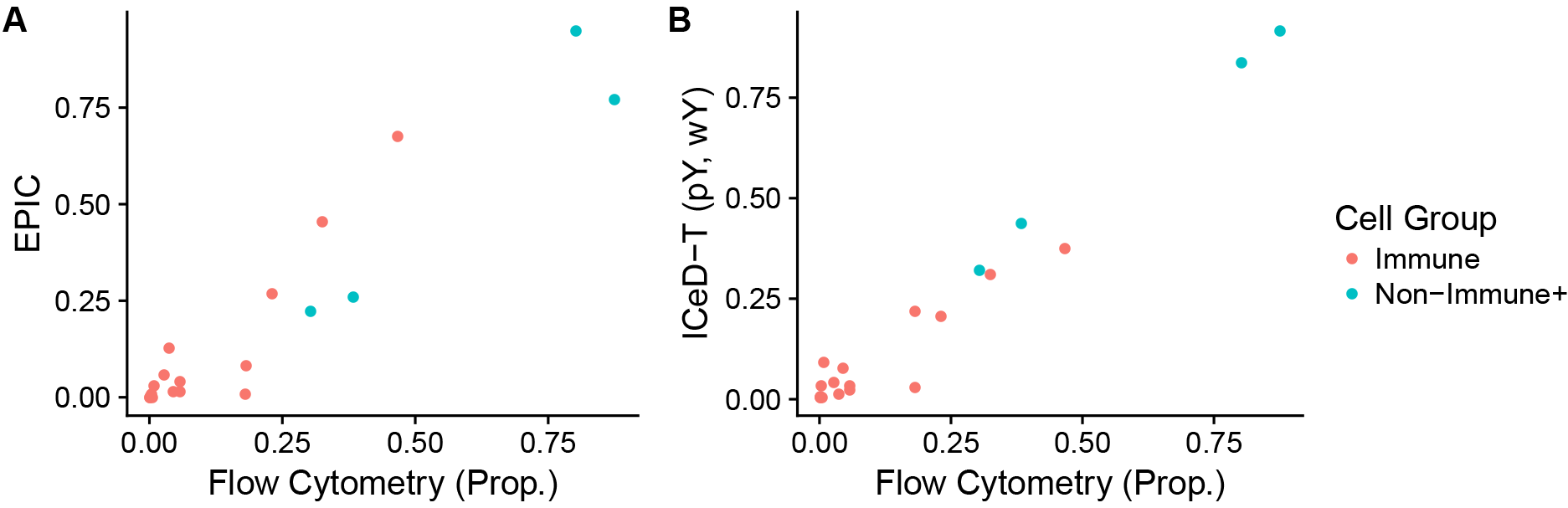
Plots of EPIC and ICeD-T model estimates against flow cytometry estimates. ICeD-T is fit using variance weights and sample purity.

**Table 2:** Sum of Squared Errors for relative immune proportions among all immune cell types. ICeD-T fits are labeled with (pX, wX) to indicate use of purity (pY=Yes and pN=No) and weight (wY=Yes and wN=No).

| Model | LAU125 | LAU1255 | LAU1314 | LAU335 |
| --- | --- | --- | --- | --- |
| CIBERSORT (LM22) | 0.12 | 0.16 | 0.003 | 0.010 |
| CIBERSORT (TRef) | 0.32 | 0.10 | 0.021 | 0.095 |
| EPIC | 0.86 | 0.15 | 0.066 | 0.013 |
| ICeD-T (pN, wN) | 1.03 | 0.10 | 0.042 | 0.003 |
| ICeD-T (pN, wY) | 1.07 | 0.14 | 0.005 | 0.004 |
| ICeD-T (pY, wN) | 0.85 | 0.08 | 0.039 | 0.008 |
| ICeD-T (pY, wY) | 0.85 | 0.14 | 0.020 | 0.002 |

We also evaluated the performance of CIBERSORT versus the flow cytometry estimates. Compared with other methods, CIBERSORT has comparable or less accurate estimates of cell type proportions in three subjects, but much better performance than the other methods in subject LAU125 (Table 2). Based on flow cytometry estimates, this subject has somewhat unexpected immune cell proportion: almost entirely B-cells. All methods perform much worse in this subject than other subjects, with larger sum squared errors. CIBERSORT has relatively better performance for this challenging subject could be due to a combination of its objective function and use of LM22 reference matrix. CIBERSORT’s performance becomes worse when using TRef instead of LM22 as reference matrix, though it still has much smaller sum squared error than EPIC and ICeD-T.

#### 3.4 Application to anti-PD-1 Immunotherapy Data

Finally, we use ICeD-T, CIBERSORT, and EPIC to analyze an RNA-seq dataset from bulk tumor samples of melanoma patients [22]. The RNA-seq data are available in 28 patients before treatment with pembrolizumab. We seek to predict treatment response (Complete Response, Partial Response, or Non-response) using CD8+ cell type composition estimated by each of the three methods.

Fastq files of RNA-seq data were downloaded from NCBI Sequence Read Archive, mapped to human genome (hg38) and the number of RNA-seq fragment per gene were counted. Then such counts were normalized by TPM. We ran EPIC and ICeD-T using the TRef reference gene expression data. ICeD-T was fit without using tumor purity as this information was not available. CIBERSORT was fit using LM22 reference matrix. Abundance estimates across each method are corrected using EPIC’s cell type size factors. In addition, to ensure comparability across all methods, immune cell proportions are renormalized so that their summation equals to 1.

Differences in relative CD8+ T-cell abundance across response categories was assessed using a Jonckheere-Terpstra test for trended differences. The Jonckheere-Terpstra test can be considered as an extension of non-parametric ANOVA tests (e.g. Kruskal-Wallis) to allow greater power to detect ordered population differences [23]. Previous studies have shown that those cancer patients with more CD8+ T cells within tumor microenviroment are more likely to respond to anti-PD-1 treatment [24]. Thus, as one moves across response categories from most to least responsive to therapy, one would expect to see a decrease in CD8+ T cell abundance.

**Figure 6:**
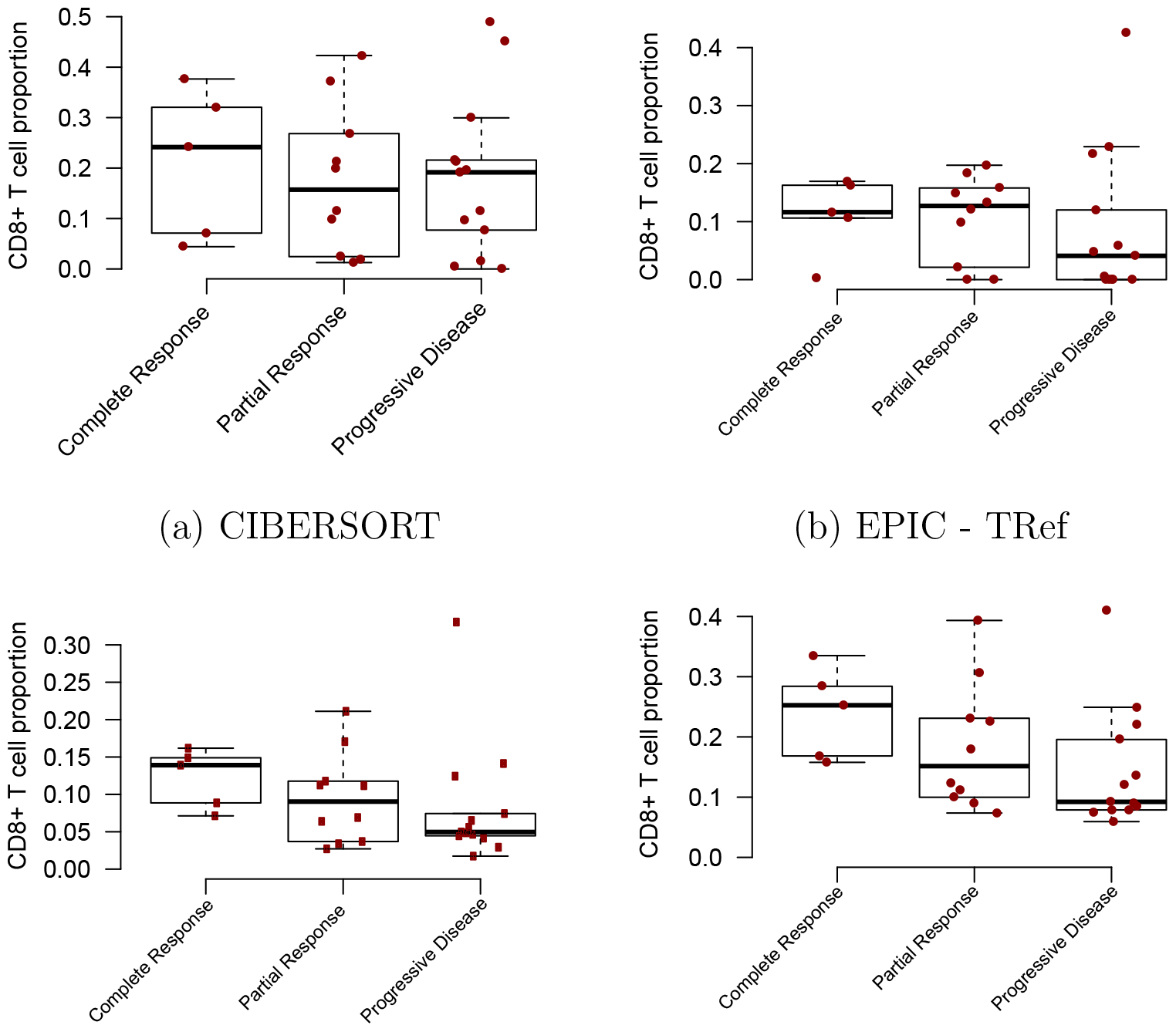
Comparison of model fits to PD-1 Immunotherapy Data

CIBERSORT and EPIC capture the expected relationship between CD8+ T cell pro-portion and immunotherapy response to some extent, but have trouble in separating the members of at least two groups. For CIBERSORT, individuals in the partial response group behave similarly to those in the progressive disease group. For EPIC, individuals in the complete response group behave similarly to those who exhibited partial response. The Jonckheere-Terpstra tests provide numerical confirmation of these difficulties as the tests are not significant, with p-values for CIBERSORT and EPIC being 0.30 and 0.14, respectively.

ICeD-T, on the other hand, provides clear visual distinction between these three groups, and shows less CD8+ T cells for those who do not respond to anti-PD-1 treatment. This relationship is reinforced through consideration of the significant Jonckheere-Terpstra test (p=0.038). Introduction of variance weights further separates these categories (p=0.017). Cell type proportions estimates by either versions of ICeD-T have higher within group similarities than either CIBERSORT or EPIC.

### 4 Discussion

In this paper, we have outlined ICeD-T, a novel statistical method for immune cell expression deconvolution within tumor tissues. ICeD-T utilizes the variance stabilizing properties of the log-transformation while simultaneously controlling for aberrant gene behavior within the tumor tissue. In addition, ICeD-T incorporates a variance weighting structure which diminishes the impact of highly variable genes on abundance estimation. Optionally, ICeD-T can refine cell type abundance estimation through use of tumor purity information, if available.

We have demonstrated that ICeD-T is an accurate model in both simulated and real datasets. The robustness of ICeD-T to misbehaved genes and its ability to identify these genes was demonstrated in simulated data. ICeD-T’s accuracy was reinforced in real datasets using both microarray and RNA-seq expression where it was consistently a top performer compared with other methods. We applied ICeD-T to study the relation between CD8+ T cell proportion and response to anti-PD-1 immunotherapy and found significant associations between CD8+ T cell proportions and patients’ response to immunotherapy.

There is room to further improve the performance of ICeD-T. One direction is to refine the reference matrix of cell type-specific gene expression. In this paper, we have adopted the reference gene expression matrix (TRef) used by EPIC. TRef was constructed using single cell RNA-seq (scRNA-seq) data from melanoma cancer samples. Cell type-specific expression was estimated by pooling cells of the same cell types, identified by clustering method. However, some technical limitations of scRNA-seq, such as dropout (expression of many genes were measured at 0 while they may be lowly expressed) may lead biased gene expression estimates [25]. Careful examination of such effects may improve the reference matrix of cell type-specific gene expression. On the other hand, scRNA-seq is a very active research area. New data analysis techniques and new data (e.g. Human Cell Atlas [26]) may help generate higher quality data for such a reference matrix.

Another future direction to improve ICeD-T is to refine the weight for each gene. We have implemented the weight for each gene based on the maximum of cell type-specific variances. Other options that use the variances across all cell types may be more desirable. However, with limited cell type-specific gene expression data, we have not yet identified a clear choice.

The ICeD-T methodology has been implemented in an R software package.

